# Evolutionary history of Euteliidae (Lepidoptera, Noctuoidea)

**DOI:** 10.1101/2022.09.26.508383

**Authors:** Reza Zahiri, Jeremy D. Holloway, Jadranka Rota, B. Christian Schmidt, Markku J. Pellinen, Ian J. Kitching, Scott E. Miller, Niklas Wahlberg

## Abstract

The Euteliidae stand apart in the Noctuoidea as the second smallest family, but most highly specialized in terms of larval hostplant relationships, favouring plant families with high levels of resin or latex, including the family dominant in Oriental lowland forests, the Dipterocarpaceae. Given the proportionally large foliage biomass represented by dipterocarps, exploitation of this abundant resource by lepidopteran defoliators is surprisingly low, and the Euteliidae may have achieved this through pre-adaption to dipterocarp chemical defences. Therefore, to assess this, we performed a molecular phylogenetic analysis on the family Euteliidae to clarify deep divergences and elucidate evolutionary relationships at the level of the subfamily, tribe, and genus. Our dataset consists of 6.3 kbp of one mitochondrial and seven nuclear DNA loci and was analyzed using model-based phylogenetic methods, i.e., maximum likelihood and Bayesian inference. We attempted to diagnose apomorphic morphological character states for Euteliidae and each monophyletic group within the family. Additionally, the evolution of hostplant use was reconstructed and a molecular-dating approach was conducted to assess the ages of major lineages within Euteliidae. We present an updated phylogenetic hypothesis for Euteliidae consisting of two strongly supported subfamilies: Euteliinae and Stictopterinae. Within Stictopterinae there are two tribes: Stictopterini and Odontodini. Several genera (e.g., *Targalla, Paectes, Marathyssa, Eutelia*) were found to be polyphyletic and require taxonomic revision. Two new genera (*Niklastelia* and *Pellinentelia*) are described and several taxonomic changes (e.g., new combinations and new synonymies) are made. The Neotropical genus *Thyriodes*, currently included in Euteliidae, is found to be associated with Erebinae (Erebidae). The divergence time estimate for the split between the Euteliidae and Noctuidae is at 53 my, and the Euteliidae split into the subfamilies Euteliinae and Stictopterinae at 42 my. In Stictopterinae, the tribes Stictopterini and Odontodini split at 31 my and the Euteliinae began a much more complex diversification at 34 my. Ancestral hostplant reconstruction identified Malpighiales (e.g., Clusiaceae) as the ancestral larval hostplant order for the family Euteliidae. The ancestors of Stictopterinae also appear to have been Malpighiales feeders, but then split into exclusive specialisms on Malvales (Odontodini) and Malpighiales (Stictopterini) hostplants. Larvae of Stictopterini appear to be restricted to Clusiaceae, apart from a few records from Dipterocarpaceae. In Euteliinae, Anacardiaceae are predominant as larval hosts. Thus, all hosts in the family are lactiferous, possibly providing some degree of pre-adaptation for exploiting Dipterocarpaceae.

## Introduction

The family Euteliidae is one of the four major lineages of the quadrifid noctuoids recognized in Zahiri et al. (2011), being a rather homogeneous and small group of 34 genera and 520 species (Nieukerken et al., 2011). Euteliidae are the only quadrifid noctuoid lineage without even a preliminary molecular phylogeny. The global diversity of euteliid moths is significantly higher in the tropics, particularly in southeast Asia, e.g., Borneo (Holloway and Barlow, 2011). Euteliidae along with three other major lineages (i.e., Nolidae, Erebidae and Noctuidae) form a robust monophyletic assemblage, informally referred to as the quadrifid noctuoids, which was discussed in detail by Zahiri et al. (2011, 2012). However, the relationships among these four families are not fully understood yet, even though collectively the group has very strong support. Quadrifid noctuoids, along with the trifid noctuoids (i.e., Oenosandridae and Notodontidae), comprise the superfamily Noctuoidea, which contains the largest radiation of Lepidoptera (moths and butterflies).

The family-level status of Euteliidae was first proposed by Mitchell et al. (2006), but both current subfamilies of Euteliidae (i.e., Euteliinae and Stictopterinae) were treated as distinct families (i.e., Euteliidae and Stictopteridae) in their phylogenetic hypothesis. The molecular analyses of Mitchell et al. (2006) confirmed these two groups as sister-groups, and in turn placed them as sister to their LAQ clade (i.e., Lymantriidae, Arctiidae, and Quadrifine Noctuoidea) within which the rest of the Erebidae groups were embedded. The close relationship between Euteliinae and Stictopterinae has also been demonstrated by Kitching (1987) and Holloway (1985) based on morphological characters. Kitching (1987) proposed 11 unique, unreversed synapomorphies uniting the two (with nine apomorphies lost in one of the included genera). Synapomoprohies uniting Euteliinae + Sticopterinae include a reduced female frenulum; a clypeofrons scaled only at the edge; weak hindtarsal spining; hindlegs shorter than midlegs; a small oval plate in the ductus ejaculatorius; ovipositor lobes (anal papillae) with inner surfaces facing at least partially posteriorly. The close relationship of Euteliinae + Stictopterinae was also discussed by Kitching and Rawlins (1998), who noted that the counter-tympanal hood has a unique double structure, although differently modified in each group. Lastly, Fibiger and Lafontaine (2005) also recognized the two taxa as sister-groups, placing them as subfamilies within their concept of Erebidae. Holloway (2011) concluded that it is more informative to treat them as distinct subfamilies within one family, for which Euteliidae is the oldest name. This arrangement was fully supported by the molecular analysis of Zahiri et al. (2011).

Adult characteristics of Euteliinae include the shape of the forewing, which is either elongate or squarish, and a hindwing that usually has a subtornal mark, often incorporated in a broad, dark border contrasting with a paler ground colour (Holloway, 1985). Lateral scale tufts at the apex of the male abdomen often give it a squared-off appearance. The male antennae are characteristically broadly bipectinate over the basal half or two-thirds, onward to the apex narrowly so or only ciliate; the antennae of females are usually filiform though occasionally partially bipectinate as in the male (Holloway, 1985). Partially bipectinate antennae of this form may be unique to Euteliinae within the quadrifid Noctuoidea but are paralleled in other families such as Notodontidae, Cossidae, and Limacodidae. The cryptic resting posture of adults of the subfamily is unusual with an upcurved abdomen that may be facilitated by musculature attached to the curved flanges of the ventral sternite at the base of the abdomen (Holloway, 1985). These flanges are distinctive and may serve to stiffen the sclerite and its development from a groove to a flange and could be a further adaptation associated with the characteristic resting posture of adults. Larval traits of Euteliidae include anterior prolegs that are not reduced, or only slightly reduced; short setae; a moderately elongate parallel-sided spinneret, and narrow spiracles (Figs 1a,b).

**Fig. 1.**
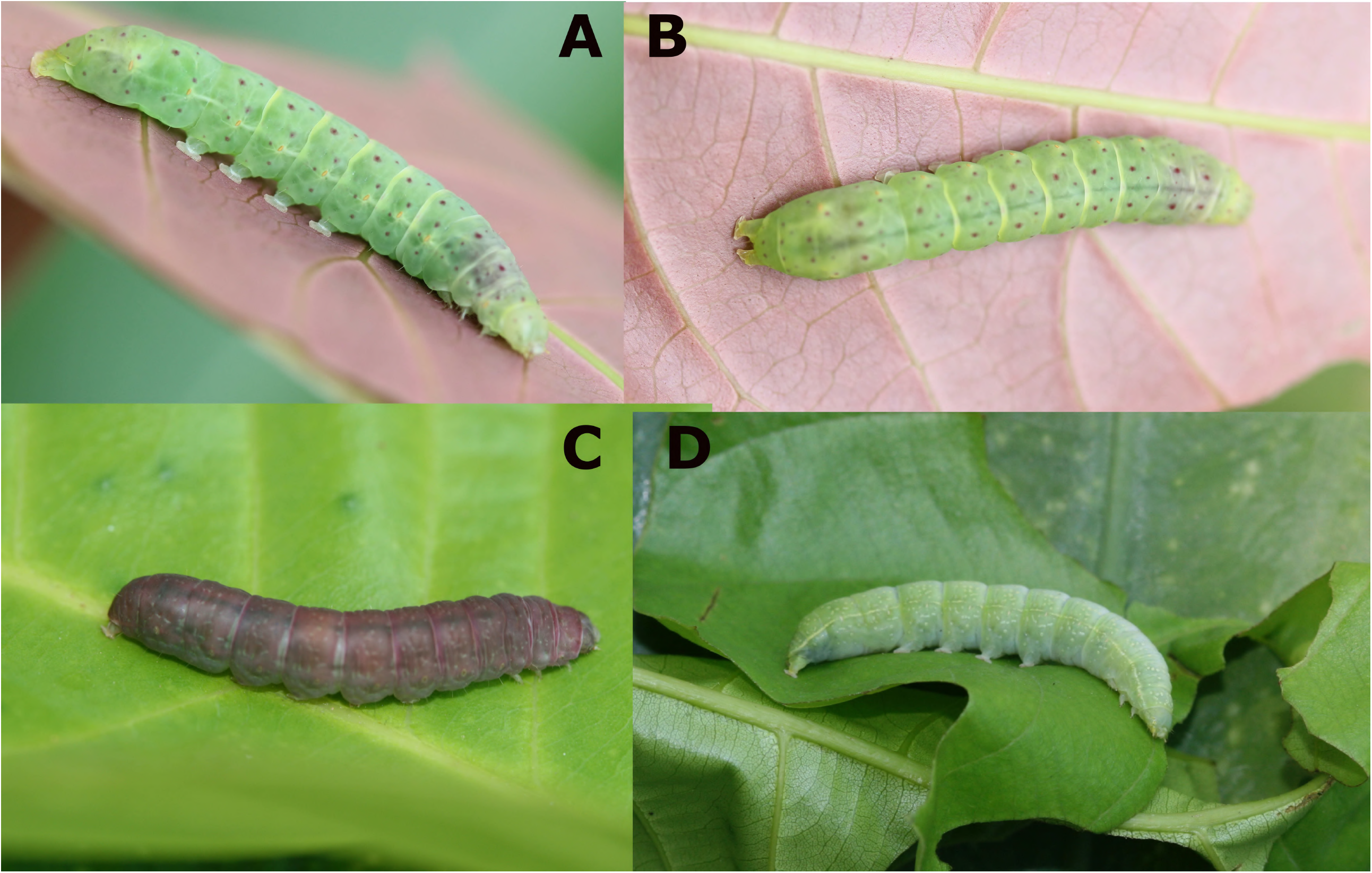
Caterpillars of Euteliidae (photographs by M. Pellinen) (A, B) 5^th^ instar of *Penicillaria jocosatrix* (Euteliinae); (C, D) 5^th^ and 4^th^ instars of *Odontodes aleuca* (Stictopterinae, Odontodini), respectively.

The prefix “sticto” in the subfamily Stictopterinae means dappled, mottled, or spotted (Holloway, 1985). The majority of species have finely complex markings on the forewings (often an etched reticulate patterning) and many genera contain species with a distinctive hyaline basal half to the hindwing, particularly *Stictoptera* Guenée, *Aegilia* Walker, and *Lophoptera* Guenée (Holloway, 1985). The antennae of both sexes are filiform, sometimes finely ciliate. The frenulum in the female is reduced to a single spine. A tuft of fine scales extends obliquely in a posterior ventral direction from anterior to the tympanum as in the Euteliinae but not to such a pronounced extent. The male eighth sternite typically bears a pair of coremata. Resting posture of Stictopterinae contrasts strikingly with the resting postures of Euteliinae. Species tend to rest with wings folded at an acute angle or, in *Stictoptera* and the more narrow-winged species, with them rolled around the body giving the appearance of a short, broken twig. Caterpillars of Stictopterinae have a complete set of four equal pairs of prolegs with uniordinal crochets, variably homoideous (*Lophoptera*), or heteroideous (*Odontodes* Guenée) (Figs 1c, d). Ventral prolegs are complete and equal, setae short and body skin smooth (Figs 1c, d).

The Euteliidae are of some minor economic importance according to records in Zhang (1994), a compendium compiled from records in the *Review of Applied Entomology* over almost a century. An indication of economic importance can be gained from the number of years in which records occur. Most Euteliinae pests defoliate fruit crops, particularly mango (*Mangifera* L.) (Anacardiaceae) with eight records for *Penicillaria jocosatrix* Guenée and a total of 12 for *Chlumetia transversa* (Walker) and *C. euthysticha* (Turner), although this genus also attacks young twigs and shoots. *Eutelia adulatrix* (Hübner) has three records from *Pistacia* L. (pistachio), also Anacardiaceae, but *E. blandiatrix* Hampson has a single record from *Altingia* Noronha (Hamamelidaceae), which is harvested for its scented resin and timber. *Targalla palliatrix* (Guenée) has two records of feeding on fruits in Myrtaceae, such as *Eugenia* L. and *Syzygium* P. Browne ex Gaertn. Fruit pests are also recorded in Stictopterinae: the *Stictoptera* species *S. bisexualis* Hampson, *S. cucullioides* Guenée and *S. grisea* Moore have five records collectively on *Garcinia* L. (mangosteen) and *Phyllanthus* L. (perfumes, oilseed, and some timber). Both subfamilies have also been recorded as forestry pests, particularly on *Shorea* Roxb. ex C.F. Gaertn. (Dipterocarpaceae), with three records for *Paectes subapicalis* (Walker) and one for *Lophoptera illucida* (Walker).

The goals of this study are first to assess the previous phylogenetic hypotheses with significantly expanded taxon sampling and then to focus on the ecology of Euteliidae, particularly the unique specialism of the early stages on a few groups of plant taxa with high levels of resin or latex in their sap and foliage that is not seen in any other group of the Macroheterocera to the same degree. In addition, we explore various morphological characters and their distributions over the molecular phylogeny.

## Material and methods

### Taxon sampling

To infer the phylogenetic hypothesis of Euteliidae, two taxon sampling strategies were adopted.

A. We inferred a backbone phylogenetic hypothesis based on a dataset of DNA sequences of eight independent protein-coding gene (PCG) regions from 183 terminal taxa, including 142 representatives of euteliid moths as the ingroup and 41 exemplars from four other families of Noctuoidea: Oenosandridae (two species), Notodontidae (three species), Noctuidae (eight species), Nolidae (four species), Erebidae (six species), and one species of Geometridae as outgroups to test the monophyly of Euteliidae (Table 1). The ingroup included representatives for all known major lineages of euteliids to date (subfamilies Euteliinae and Stictopterinae (including two tribes, Odontodini and Stictopterini)) and represents 20 of the 34 known genera of euteliids (Table 2). This selection was mainly based on the results of previous morphological and molecular works (Fibiger and Lafontaine, 2005; Lafontaine and Fibiger, 2006; Lafontaine and Schmidt, 2010; Zahiri et al., 2013b, 2013a, 2012, 2011). Cladograms were rooted with Geometridae, which represents the hypothesised sister taxon to Noctuoidea (Mutanen et al., 2010; Nieukerken et al., 2011; Regier et al., 2009).
B. To find a home for a wide range of euteliids currently labeled as “*incertae sedis*” — i.e., their placement is debated at ranks above the species level — we added DNA barcode data from 242 specimens to the dataset in A (Table 3) to produce a 425-specimen dataset of Euteliidae (384 ingroup and 41 outgroup exemplars) (Table 3). This dataset includes 26 of the 34 currently recognized genera of euteliids (Table 2). A phylogeny was then generated that highlighted potential taxonomic misplacements, unrecognized major clades, and positions for *incertae sedis* taxa. We use these results to make systematic rearrangements within the family Euteliidae.

### Gene sampling

Total genomic DNA was extracted from one or two legs, dried or freshly preserved in 96% ethanol, using the DNeasy tissue extraction kit (Qiagen, Hilden, Germany) or the NucleoSpin Tissue 250 kit (Macherey-Nagel, Düren, Germany) and following the manufacturers’ instructions. For each specimen, we sequenced eight protein-coding exons that have previously been found to be highly informative in higher-level lepidopteran phylogeny (Keegan et al., 2021, 2019; Mutanen et al., 2010; Rota et al., 2016; Wahlberg et al., 2009; Wahlberg and Wheat, 2008; Zahiri et al., 2013b, 2013a, 2012, 2011). The gene regions were single-copy from the nuclear genome: elongation factor-1α (EF-1α), ribosomal protein S5 (RpS5), carbamoylphosphate synthase domain protein (CAD), cytosolic malate dehydrogenase (MDH), glyceraldehyde-3-phosphate dehydrogenase (GAPDH), isocitrate dehydrogenase (IDH) and wingless genes, and, from the mitochondrial genome, cytochrome c oxidase subunit I (COI). We followed the protocols from Wahlberg and Wheat (2008) for DNA amplification (PCR) and pre-sequencing methods. We sequenced our PCR products at the CFIA Ottawa Plant Laboratory (Fallowfield, Ontario, CA), or sent them to either Macrogen Europe Inc. (Amsterdam, The Netherlands) or Macrogen, Seoul, Korea for Sanger sequencing. For the majority of loci, we used single forward reads, although for some fragmented PCR products we used reverse reads as well (Wilson 2012). We have deposited the sequences generated for this study in GenBank (GenBank accession numbers will be added to Table 1).

### DNA sequence data and quality controls

We inspected sequence chromatograms visually for base call errors and heterozygous loci, and produced consensus sequences in Geneious® version 10.2.5 (http://www.geneious.com, Kearse et al., 2012), BioEdit (Hall, 1999), or Mesquite version 3.2 (Maddison and Maddison, 2017). Alignment was trivial and the few insertion/deletion events that were detected were of entire codons that could be easily aligned. To minimize the risk of any confusion during the sequencing protocol and errors in alignments, we constructed neighbor-joining and Maximum Likelihood trees separately for each gene region and checked them carefully for identical sequences and other doubtful patterns. In addition, to minimize the risk of incorrect identifications, all the specimens with COI sequences were cross-checked with their DNA barcodes in BOLD (Barcode of Life Data System, http://www.boldsystems.org) (Ratnasingham and Hebert, 2007), where reference specimens were available for many of the species used in this study.

### Phylogenetic analyses

The aligned DNA sequences were added to a VoSeq version 1.7.4 database (Peña and Malm, 2012) from which we generated aligned datasets. The gene regions were explored using model-based phylogenetic approaches including Maximum Likelihood (ML) and Bayesian Inference (BI) methods. The dataset was analyzed using the maximum likelihood framework in the program IQ-TREE V1.6.12 (Nguyen et al., 2015). The best-fitting substitution models and the optimal number of partitions were selected by ModelFinder (Kalyaanamoorthy et al., 2017). Support for nodes was evaluated with 1000 ultrafast bootstrap (UFBoot2) approximations (Hoang et al., 2018), and SH-like approximate likelihood ratio test (Lanfear et al., 2012). SH-Like ≥80 and UFBoot2 ≥ 95 values indicate well-supported clades. To reduce the risk of overestimating branch supports in UFBoot2 test, we implemented the -bnni option, which optimizes each bootstrap tree using a hill-climbing nearest neighbour interchange (NNI) search. BI analyses were carried out using the software MrBayes v3.1 (Ronquist et al., 2005) on the freely available Bioportal server (http://www.bioportal.uio.no). The Bayesian analyses were run independently two times for 20 million generations, with every 1000th tree sampled. The dataset was divided into two partitions: mtDNA and nDNA, as partitioning by gene resulted in poor mixing of chains and problems with convergence. We modeled the evolution of sequences according to the GTR+ Γ model independently for the two partitions using the ‘‘unlink’’ command in MrBayes. Clade robustness was estimated by posterior probabilities in MrBayes. Convergence was determined when the standard deviation of split frequencies went below 0.05 and the PSRF (Potential Scale Reduction Factor) approached 1, and both runs had properly converged to a stationary distribution after the burn-in stage (which was 1,000 sampled generations). The Interactive Tree Of Life (https://itol.embl.de) was applied for the display, manipulation and annotation of phylogenetic trees (Letunic and Bork, 2021).

### Time of divergence

The multi-locus dataset with 183 taxa (Table 1) was used to estimate dates of cladogenetic events at the family level as well as lower taxonomic groupings (e.g., subfamilies, tribes, and generic level). Times of divergence were estimated using a Bayesian lognormal relaxed clock with uncorrelated rates to each clock model as implemented in BEAST 2.6.3.0 (Bouckaert et al., 2019). The dataset was partitioned by genes and the best-fit model of evolution of sequences, according to BIC, was found using ModelFinder (Kalyaanamoorthy et al., 2017) in IQ-TREE (Nguyen et al., 2014; Trifinopoulos et al., 2016). As a result, we unlinked the partitions for the substitution models, allowing partitions to independently evolve under different substitution models (see Table S1). However, we linked the partitions for the clock and tree models (the list of best-fit substitution models per partition can be found in Table S1). The tree prior was set to a Birth-Death Model in one run, and a Yule Model in another run. Ditrysian Lepidoptera are unfortunately characterized by a lack of fossils that can be confidently assigned to extant clades (Grimaldi and Engel, 2005; Wahlberg et al., 2013), and Euteliidae are no exception. Even the fossils that are assigned to the superfamily Noctuoidea (Sohn et al., 2012), are either ambiguous (higher taxonomy assignments are mostly weakly founded), or have been identified based on partial body or single wings, or are not old enough (Last Glacial Maximum/Late Pleistocene) to inform node calibrations of Noctuoidea (Kristensen, 1998). We used the recent phylogenomic study of Kawahara et al. (2019) to apply secondary calibration points for the estimation of divergence times of the Euteliidae. We constrained the root of the superfamily Noctuoidea (69 my) with normal distributions encompassing the 95% credibility intervals estimated in Kawahara et al. (2019). We then constrained the roots of the family Erebidae and the subfamily Arctiinae to 47.8 my and 30 my, respectively, with normal distributions encompassing the 95% credibility intervals (see Dataset S11 Kawahara et al. 2019). The MCMC parameters were fixed to 150 million generations for two separate runs (tree priors set to a. Birth-Death Model and b. Yule Model) with sampling every 5,000 generations and the first 25% discarded as burn-in.

### Hostplant ancestral state reconstruction

Ancestral state reconstruction was conducted using RASP (Reconstruct Ancestral State in Phylogenies, version 4.2) (Yu et al., 2015) using the Bayesian Binary MCMC (BBM; Ronquist and Huelsenbeck 2003) for hostplant order association: (A) Sapindales, (B) Fagales, (C) Rosales, (D) Myrtales, (E) Malpighiales, (F) Malvales, (G) Fabales, (H) Laurales, (I) Saxifragales (Table 4). Hostplant information was compiled from the literature (Gardner, 1948; Holloway, 1985; Janzen and Hallwachs, 2017; Mathur, 1942; Robinson, 1974; Robinson et al., 2010; Sevastopulo, 1941; Staude et al., 2020). Higher-level classification and nomenclature of plants are based on the Angiosperm Phylogeny Group (APG) classification of the orders and families of angiosperms (The Angiosperm Phylogeny Group et al., 2016). BBM analysis was based on 10,001 trees, of which 3,000 were discarded (burn-in 30%), sampled from one run of a Bayesian inference calculated by MrBayes 3.2 as outlined above but using a reduced alignment (122 terminal leaves) to include only ingroups with reliable foodplant records (Table 4). A condensed tree was calculated and the outgroups were removed by using the tools provided by RASP.

## Results

### Phylogenetic hypothesis

Based on two model-based phylogenetic approaches (ML and BI) for the combined datasets, our results strongly support the monophyly of Euteliidae (100/1/100/1; numbers in parentheses are SH-aLRT support (%) / aBayes support / ultrafast bootstrap support (%) / Posterior Probability respectively) (Figs 2, 3a, 3b). Euteliidae consist of two strongly supported lineages, the subfamilies Euteliinae (100/1/100/1) and Stictopterinae (100/1/100/1) (Figs 3a, 3b). The optimal topologies from the two analyses are very similar, with slight variation observed mostly in the terminal branches and slightly in branch supports. The analysis of the partitioned dataset by mtDNA and nDNA genes, as well as the dataset omitting the mitochondrial COI gene, did not have a significant impact on the tree topology and node supports.

**Fig. 2.**
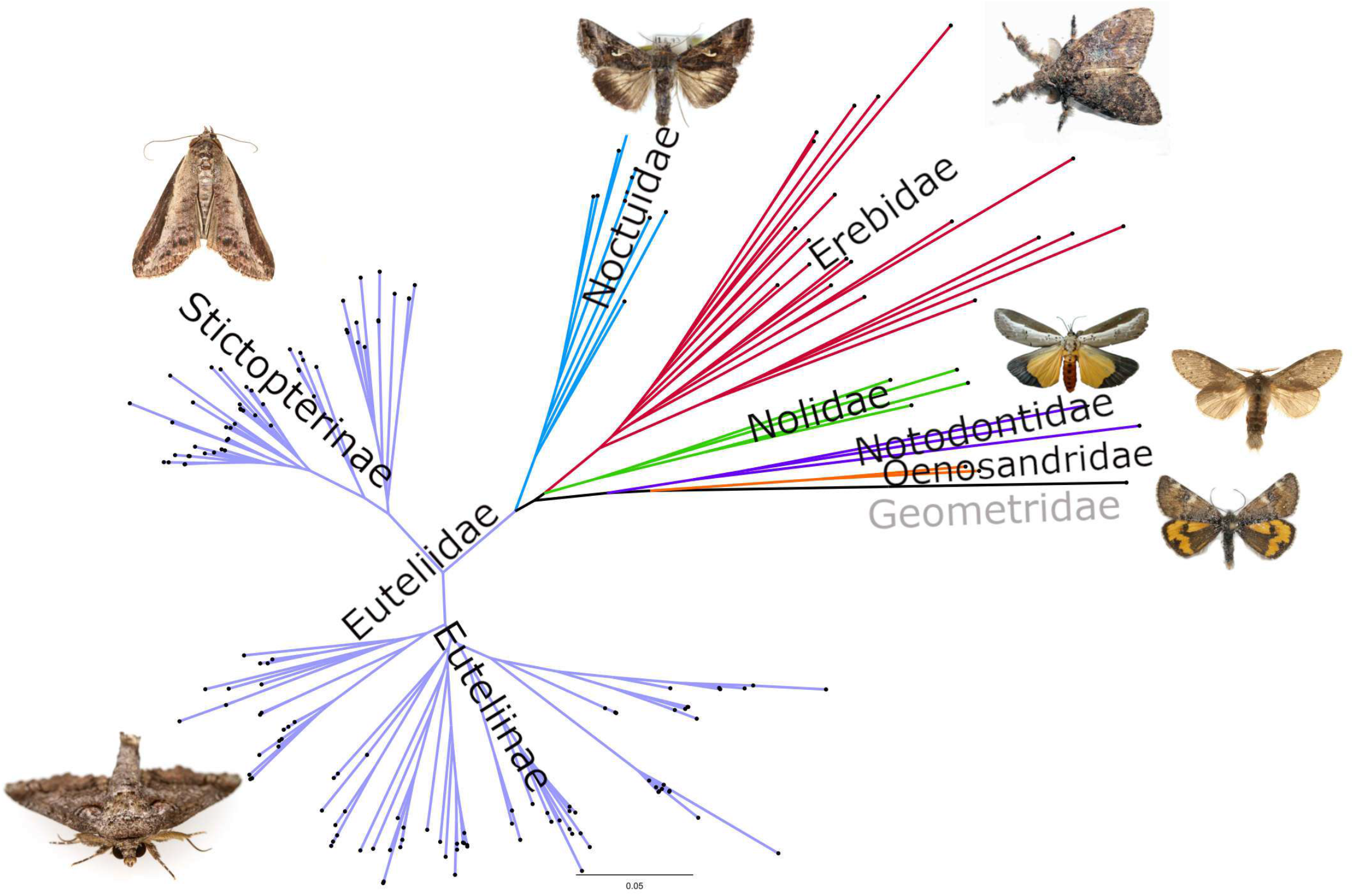
Phylogenetic hypothesis of Noctuoidea based on Maximum Likelihood and Bayesian inference analyses. Clades representing major lineages (families) are coloured.

**Fig. 3a & b.**
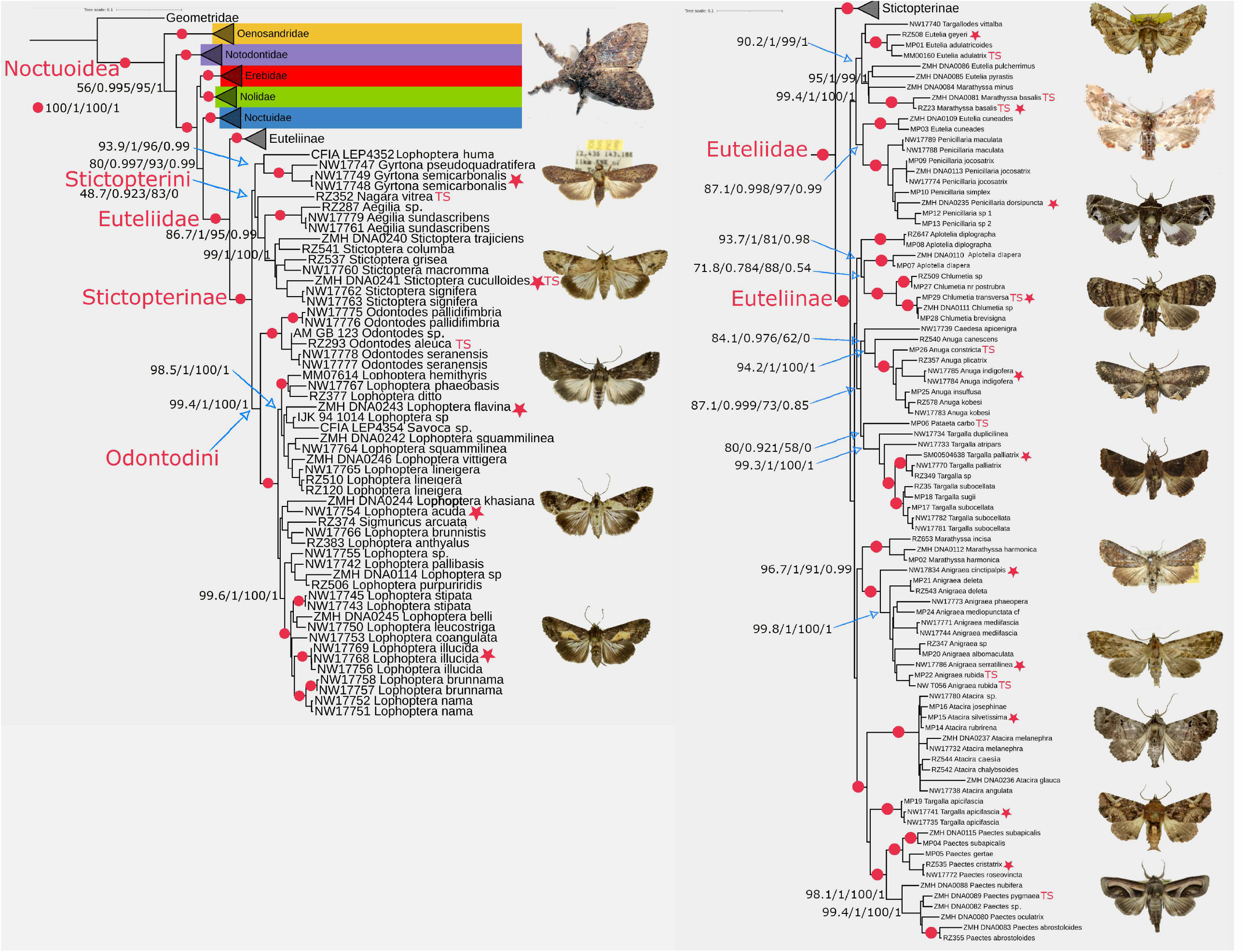
Phylogenetic hypothesis of Euteliidae and out-groups based on Maximum Likelihood and Bayesian inference analyses. Clades representing outgroups are collapsed and coloured. Support values shown next to the branches are SH-aLRT support (%) / aBayes support / ultrafast bootstrap support (%) / posterior probabilities, respectively. The names of the moths in the figure are associated with the terminal leaves marked with red stars. **3a**. Out-groups and Stictopterinae; **3b**. Euteliinae

Relationships amongst the four families within the quadrifid Noctuoidea (i.e., Euteliidae, Noctuidae, Erebidae, and Nolidae) are not clear, although they form a strongly supported (100/1/100/1) monophyletic group (Figs 2, 3a, 3b). Euteliidae are sister to Noctuidae, and Nolidae are sister to Erebidae in both ML and BI analyses with moderate to weak supports (80/0.997/93/0.99; 59.5/0.95/94/0.99 respectively, Figs 2, 3a, 3b).

#### Subfamily Stictopterinae

Stictopterinae are recovered as a strongly-supported monophyletic group including two moderately supported major lineages: one comprised of the genera ((*Stictoptera, Aegilia*), *Nagara* Walker, *Gyrtona* Walker) (48.7/0.923/83/0.9) (tribe Stictopterini), and the other lineage (tribe Odontodini) (99.4/1/100/1) includes ((*Odontodes*), (*Lophoptera*/*Sigmuncus* Holloway/*Savoca* Walker)) (Fig. 3a). *Lophoptera huma* (Swinhoe) is nested as sister to *Gyrtona* clade.

#### Subfamily Euteliinae

Our analysis recognized three strongly supported generic groupings, together with a weakly supported lineage (87.1/0.999/73/0.85) represented by the six genera *Aplotelia* Warren (TS) (including *Aplotelia diapera* (Hampson) see below); *Chlumetia* Walker (TS); *Caedesa* Walker, *Pataeta* Walker (TS), *Anuga* Guenée (TS), and *Targalla* Walker, within the subfamily Euteliinae (Fig. 3b). The first clade (99.4/1/100/1) includes the genera *Eutelia* Hübner (TS), *Targallodes* Holland, *Marathyssa* Walker (TS) and *Penicillaria* Guenée. The second clade (96.7/1/97/0.99) comprises the monophyletic genus *Anigraea* Walker (TS) and the Oriental *Marathyssa* (i.e., *M. harmonica* (Hampson) and *M. incisa* Kobes). And finally, the third clade (100/1/100/1) contains the Nearctic and Old World *Paectes* Hübner (TS) + *Callingura* Butler, monophyletic genus *Atacira* Swinhoe (TS), and *Targalla apicifascia* (Hampson) (Fig. 3b).

Within the first clade, the true *Eutelia-*group (including the TS) is associated with the Afro-Oriental genus *Targallodes* with weak support (43.6.7/0.9/92/0.7). This clade (*Eutelia* + *Targallodes*) is associated with a monophyletic group (95/1/99/1) that contains the Nearctic *Marathyssa* (including the TS) and *Eutelia* (*E. pyrastis* Hampson and *E. pulcherrimus* (Grote)), forming a moderately-supported lineage (90.2/1/99/1) (Fig. 3b). *Eutelia* is polyphyletic, with the Old World type species (*E. adulatrix*) associated with *E. geyeri* (Felder and Rogenhofer) (TL: Japan) and *E. adulatricoides* (Mell) (TL: China), whereas two Nearctic species (*E. pyrastis* and *E. pulcherrimus*) are allied with the Nearctic genus *Marathyssa*, and finally, two exemplars of the Asian species, *E. cuneades* (Draudt) are associated with the Indo-Australian genus *Penicillaria* with moderate support (87.1/0.998/97/0.99). *E. cuneades* was originally placed in the genus *Marathyssa* but Sugi (2000) transferred it to *Eutelia* as a close relative to *E. favillatrixoides* Poole. Moreover, *Marathyssa* is polyphyletic, with the New World type-species (*M. basalis* Walker) positioned with another Nearctic species (*M. minus* Dyar), along with two Nearctic species (*E. pyrastis* and *E. pulcherrimus*), forming a strongly supported clade (95.1/1/99/1), and the other two Oriental species *M. harmonica* and *M. incisa* are placed with strong support (96.7/1/91/0.99) as sister to the monophyletic assemblage that includes the type-species of the Indo-Australian *Anigraea* and its associated species with strong support (100/1/100/1) (Fig. 3b).

The Indo-Australian genus *Penicillaria* is supported as monophyletic (Fig. 3b). Our results suggest paraphyly of the Indo-Australian genus *Aplotelia* (Fig. 3b) with *A. diapera* grouped as sister to monophyletic Afro-Oriental genus *Chlumetia* (71.8/0.784/88/0.54) comprising the type species (*C. transversa* Walker), and *A. diplographa* Hampson placed as a sister to *A. diapera* + *Chlumetia* (93.7/1/81/0.98) (Fig. 3b).

The Indo-Australian genus *Targalla* was found to be polyphyletic since *T. apicifascia* is not associated with other *Targalla* species (seven species) that form a monophyletic group (99.3/1/100/1). The latter clade arises as a sister to the type species of the Old World Tropical genus *Pataeta* Walker (Fig. 3b) with weak supports (80/0.921/58/0). The Southeast Asian genus *Caedesa* is paired with the *Anuga* clade (94.2/1/100/1) —including the TS—with weak support (94.1/0.967/62/0) (Fig. 3b).

Multi-locus phylogenetic analyses showed a deep evolutionary relationship between the New World *Paectes* clade including the TS, *P. pygmaea* Hübner (98.1/1/100/1), and the Old World *Paectes* (*Callingura*) (100/1/100/1) with strong support (100/1/100/1). This pair (the Nearctic *Paectes* + Old World *Paectes* (*Callingura*) is sister to *Targalla apicifascia* with no support (Fig. 3b). Similarly, this triplet is sister to the monophyletic assemblage of nine species of *Atacira* (including the TS) with strong support (100/1/100/1).

### Backbone phylogeny and barcode data

Our results based on ML analysis of 435-specimen dataset of Euteliidae and outgroups with nearly 43% of representative species included in the multigene backbone and the remaining 57% included only by mitochondrial COI barcode data, provide strong support (BS = 99) for the monophyly of Euteliidae comprising two strongly supported clades (i.e., subfamilies): Euteliinae (BS = 98) and Stictopterinae (BS = 98) (Fig. S1).

Our molecular results exclude the Neotropical *Thyriodes* Guenée from Euteliidae, instead placing it among the *Zale* Hübner and *Catocala* Schrank lineages of Erebidae (subfamily Erebinae) (Fig. 4), with strong support (BS = 100). The rest of the Euteliidae exemplars represented by barcode data grouped are within the Euteliidae clade. Within Stictopterinae, there are two major lineages (i.e., tribes) with good support: Stictopterini and Odontodini. The tribe Stictopterini contains monophyletic genera *Stictoptera, Nagara, Aegilia* and *Gyrtona* (Fig. 4, S1). *Gyrtona* appears to be sister to the rest of the genera within Stictopterini (Fig. 4). The tribe Odontodini comprises monophyletic genera *Odontodes* and *Lophoptera* (Fig. 4). *Sigmuncus* is placed in a monophyletic group as sister to some Bornean species within a larger species-group of *Lophoptera* (Fig. 4, S1). *Diascoides* Holloway, with a single species and only barcode data, is grouped between *Lophoptera* and *Odontodes* (Fig. 4). Finally, representatives of *Savoca* are placed in a clade as sister to Bornean *Lophoptera univalva* Holloway within a larger group of *Lophoptera* species (Fig. 4, S1).

**Fig. 4.**
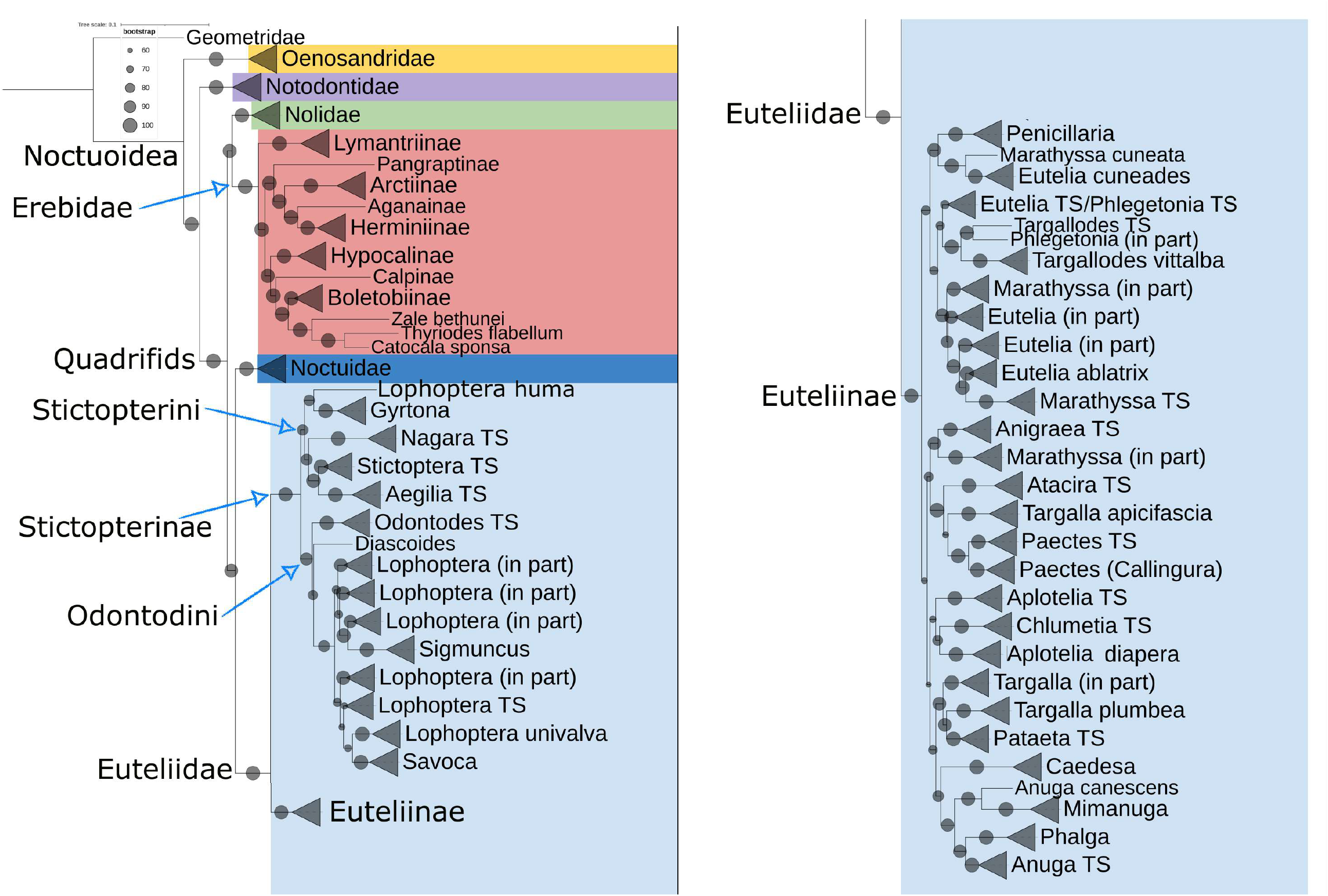
Phylogenetic hypothesis of Euteliidae and outgroups inferred by Maximum Likelihood analysis. The MLtree contains 425 terminal taxa in total: 183 with multi-locus sequence data and 252 terminals represented only by mitochondrial COI barcode data. Bootstrap values (please see legend) are displayed as red circles (five various sizes) on the tree branches. The scale bar shows the estimated number of substitutions per site. Monophyletic generic clades are collapsed.

Within the subfamily Euteliinae, *Penicillaria* is recovered as a monophyletic genus (BS = 100) and is placed as a sister (BS = 97) to a strongly supported clade (BS = 100) that contains *Eutelia cuneades* from Thailand and *Marathyssa cuneata* (Saalmüller) from South Africa (Fig. 4, S1). This pair is weakly (BS = 80) associated with a clade with weak support (BS = 77) consisting of ((*Targallodes*/*Phlegetonia* Guenée), (*Eutelia* TS/*Phlegetonia* TS)), (*Marathyssa* TS/*Eutelia*) (Fig. 4, S1).

*Aplotelia* (contains the TS) is recovered as a monophyletic group and is placed, with weak support, as sister (BS = 69) to a poorly supported clade (BS = 83) that contains the monophyletic Afro-Oriental genus *Chlumetia* comprising the type species and a South Asian species (*diapera*) of the genus *Aplotelia* (Fig. 4, S1). Two exemplars of the Old World tropical genus *Mimanuga* Warren from PNG + a singleton Bornean *Anuga* (see below) are nested within a larger clade with good support (BS = 94) that contains the true *Anuga* clade (BS = 100) and *Phalga* Moore (Fig. 4, S1). This clade (*Anuga* TS + *Phalga* TS + *Mimanuga*) is placed as a sister to *Caedesa* with moderate support (BS = 87). The Bornean *Anuga canescens* appears to be more associated with *Mimanuga* than with the core *Anuga* clade (Fig. 4, S1).

*Targalla* appears to be polyphyletic, as the Indomalayan *Targalla apicifascia* is placed within a strongly supported clade (BS = 100) that contains *Atacira* and *Paectes*, as sister to *Paectes* with weak support (BS = 67), *Targalla plumbea* (Walker) is placed as a sister lineage to *Pataeta* (contains TS), with good support (BS = 96), and a large group of *Targalla* species with strong support (BS = 100) is placed as sister to *Pataeta*+*Targalla plumbea* (Fig. 4, S1).

Old World *Paectes* forms a strongly supported monophyletic group that includes *Callingura* Butler (Fig. 4, S1). This assemblage (*Paectes*/*Callingura*) is placed as a sister to Indomalayan *Targalla apicifascia*, but with only weak support (BS = 67). This weakly supported pairing is a sister to the monophyletic genus *Atacira* (TS) (BS = 100), with a strong support value (BS = 100) (Fig. 4, S1). This triplet is sister to another well-supported clade (BS = 95) including *Anigraea* (TS) + *Marathyssa incisa*/*harmonica*, but this relationship is poorly supported (BS = 80) (Fig. 4, S1).

### Time of divergence

The chronogram (Fig. 5) obtained through the Bayesian dating analyses allows us to infer a median age of 41.87 my (mid-Eocene) for the crown clade of Euteliidae, with 95% highest posterior densities (HPD) of 34.7–49.43 my. Euteliidae diverged from the common ancestor with Noctuidae around 53.10 my (95% HPD: 45.04–61.28 my). Our analysis of divergence time suggests that the two major lineages of Euteliidae began diversifying in the early Oligocene some 31–34 my (Fig. 5). Within Stictopterinae, beginning with the bifurcation of Stictopterini from its sister group, Odontodini, various clades appear to have diverged throughout the late Oligocene (Fig. 5). Within Euteliinae, the diversification of major groups appears to be approximately the same age as the Stictopterinae tribes, i.e., about 25–28 my (Fig. 5).

**Fig. 5.**
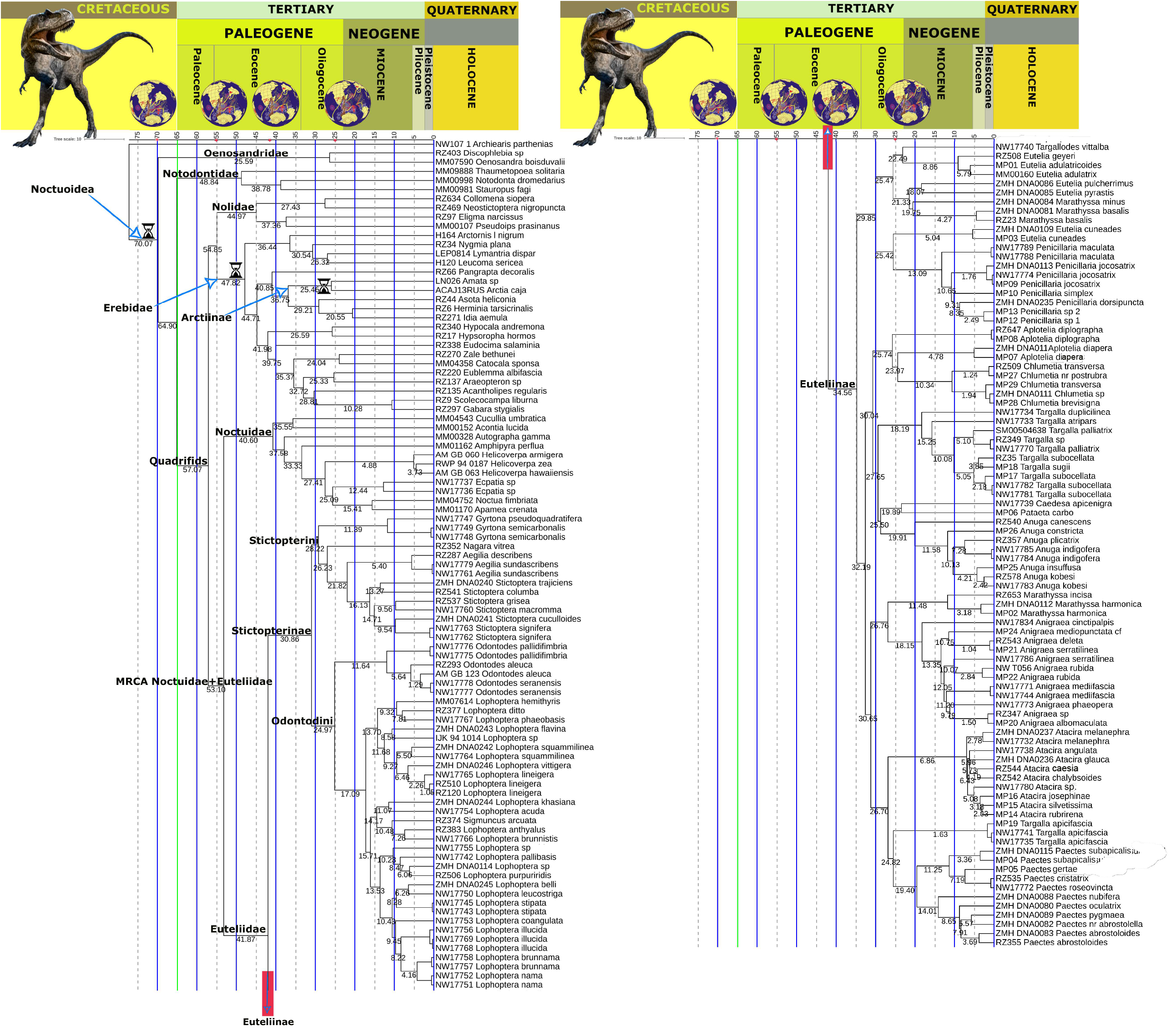
Maximum credibility tree with median ages (my) from the Bayesian uncorrelated uniform analysis under BEAST for the superfamily Noctuoidea. A five my-timescale is placed at the top of the chronogram and spans the Cretaceous to the Holocene. Black hourglasses at the corresponding nodes indicate the calibration points used to ultrametrize the topology. For the groups of major interest, the name is given above the branch leading to the group.

### Reconstruction of Ancestral Character States (Hostplant Associations)

The hostplant order Malpighiales is recovered as the most probable ancestral state for the crown of Euteliidae (Fig. 6). The ancestors of the subfamily Stictopterinae appear to have been Malpighiales feeders (69.49%), the same as the ancestors of the tribe Stictopterini (96.33%) (Fig. 6). However, our phylogenetic reconstructions suggest that there has been a split between Malvales+Malpighiales (71.99%) / Malvales (25.89%) (Odontodini) and Malpighiales (96.33%) (Stictopterini) specializations (Fig. 6). Within Euteliinae, most of the lineages leading to genera were predominantly feeding on Sapindales (94.63%). Our results suggest that there have been multiple shifts at the generic level from Malpighiales to new hostplant orders, e.g., to Laurales, Malvales, Fagales, Fabales, and Myrtales (*Paectes*), to Sapindales (*Chlumetia, Atacira, Penicillaria, Aplotelia*), to Sapindales, Fabales, and Fagales (*Eutelia, Marathyssa*), to Fabales and Sapindales (*Pataeta*), to Myrtales (*Targalla*), to Sapindales, Fagales, Fabales and Myrtales (*Anigraea*), to Sapindales, and Fabales (*Anuga*) (Fig. 6).

**Fig. 6.**
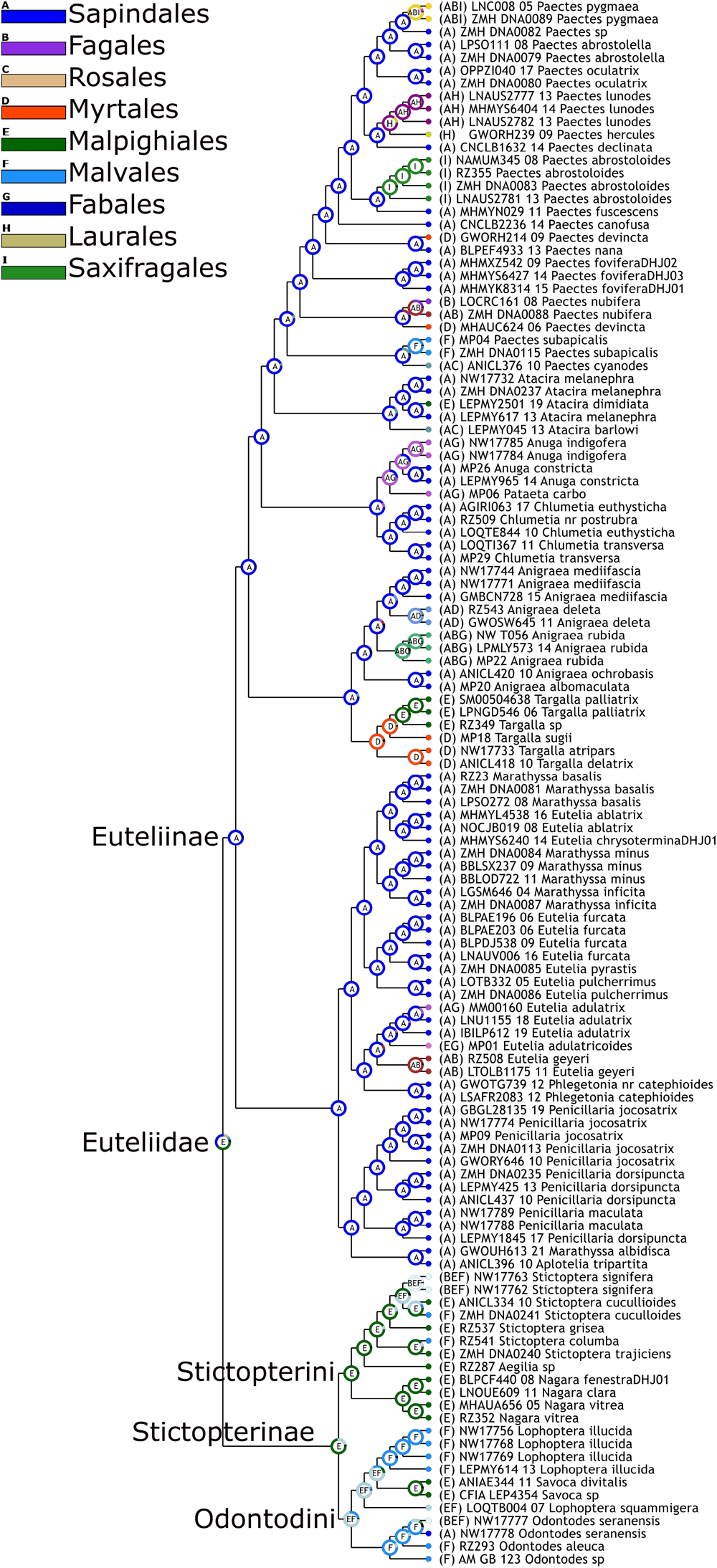
Ancestral trait reconstruction for Euteliidae hostplant Orders using RASP 4.2. Stochastic character mapping with 100,000 replicates. Most Likely States (MLS) at each node are displayed in the centre of pie charts. Hostplant Orders are: A) Sapindales, B) Fagales, C) Rosales, D) Myrtales, E) Malpighiales, F) Malvales, G) Fabales, H) Laurales, I) Saxifragales.

## Discussion

### Systematics and taxonomy

The Euteliidae lineage is undoubtedly a distinct, well-defined group of quadrifid noctuoids, deserving family status based on the multi-locus phylogenies (Zahiri et al., 2013b, 2013a, 2012, 2011) and morphology and biology (H olloway 1985, 2011; Kitching 1987; Lafontaine and Schmidt 2010). The two lineages within Euteliidae, Stictopterinae, and Euteliinae, are strongly supported as monophyletic and as sister groups (Figs 2, 3). The relationship of Euteliidae to the three remaining quadrifid Noctuoidea families has been unclear. In this study, Euteliidae are associated with Noctuidae, and Nolidae are sister to Erebidae in both ML and BI analyses (Figs 2, 3). This is in contrast to the results of Mitchell et al. (2006) where both Euteliinae *s*.*s*. + Stictopterinae *s*.*s*. grouped as the sister group to Erebidae. However, the sister group relationship between Euteliidae and Noctuidae was also recovered by Zahiri et al. (2011, 2012) and a study based on the entire mitochondrial genome also placed Euteliidae (with one exemplar species) as the sister to Noctuidae (Yang et al., 2015). More recently, a new study based on approximately 2,600 loci using hybridization capture also found Euteliidae as a sister group to Noctuidae (Mayer et al., 2021). These results strongly corroborate a strongly supported sister relationship between Euteliidae and Noctuidae.

The close association of the two current subfamilies (Euteliinae and Stictopterinae) within Euteliidae had already been examined and suggested by Richards (1933), Holloway (1985), and Kitching (1987). Numerous synapomorphies have been previously proposed to unite Euteliinae + Stictoperinae, including reduced female frenulum, modified basiconic sensilla on the proboscis, presence of a small oval plate in the ductus ejaculatorius, inner surface of anal papillae directed posteriorly, and the counter-tympanal hood consisting of a unique double structure (Holloway, 1985; Kitching, 1987; Kitching and Rawlins, 1998; Richards, 1933). Our morphological examinations supported the above mentioned symapomorphies being shared within the family members of Euteliidae.

Within the subfamily Euteliinae there is strong support for the relationships among the genera *Eutelia* (represented by the TS), *Targallodes, Marathyssa* (represented by the TS), *Phlegetonia* (represented by the TS), and *Penicillaria* (Figs 3, 4). Our phylogenetic hypothesis confirmed polyphyly of the genus *Marathyssa*, requiring the two Oriental species (*M. harmonica* and *M. incisa*) from Thailand and Borneo, which are more closely related to the Oriental genus *Anigraea*, to be examined morphologically more critically (Figs 3, 4). Our phylogeny also revealed a deep split within the *Paectes* + *Callingura* clade (Fig. 4), as the New World species (including the TS) are united in a monophyletic assemblage that appears to be sister to Oriental *Paectes* + *Callingura* species (*P. roseovincta* Warren, *P. cristatrix* (Guenée), *C. gertae* Kobes, *C. subapicalis* (Walker)). Kobes (1994–2008) revived *Callingura* as a full genus (the name *Callingura* is available as a subgenus for the Indo-Australian taxa, Holloway 1985), distinct from the New World *Paectes* Hübner. However, Kobes did not revise *Callingura* but simply preferred to use it for the Oriental taxa, whereas Holloway felt the need in his Moths of Borneo volume 14 (1985) to recognize the monophyly of *Paectes* as a whole to retain the New World/Oriental connection. Afterward, Kononenko and Pinratana (2013) said they followed Holloway (1985, 2011) and treated *Callingura* as a synonym of *Paectes*, e.g., “*Paectes gertae* (Kobes, 2008)”. In fact, they go as far as saying “NOTE: The species [*gerthae*], described in *Callingura* (Kobes, 2008) belongs to the genus *Paectes* (Holloway, pers. comm.).” So unless someone reinstated *Callingura* as a (sub)genus after that, or we ignore the non-explicit synonymy in Moths of Thailand, this will be a **stat. rev**. here. As a result, those Oriental species that are currently treated in the genus *Paectes* (i.e., *P. roseovincta, P. cristatrix*) should be included in the subgenus *Paectes* (*Callingura*), and those New World species that are currently treated in the genus *Paectes* (Table 5) should be included in the subgenus *Paectes* (*Paectes*). Our divergence time analysis suggests the cladogenesis began to occur around 19.40 my resulting in an evolutionary splitting into two lineages of New World species and Oriental species of *Paectes* (Figs. 3b, 4, 5).

The Asian genus *Targalla* is found to be polyphyletic since *T. apicifascia* is not associated with other *Targalla* (including the TS), as first suggested by Holloway (1985). It clearly requires a new genus, more closely associated with the *Paectes* clade (*Paectes* (*Paectes*) and *Paectes* (*Callingura*) (Figs 3b, 4, S1), which we describe below. *Eutelia* is recovered as polyphyletic, with the Nearctic species and *E. cuneades* from Thailand falling outside the true *Eutelia* group (Figs 3b, 4, S1). Two Nearctic species (*E. pyrastis* and *E. pulcherrimus*) are grouped within the Nearctic *Marathyssa* (including the TS), and *E. cuneades* from Thailand is closely associated with *Penicillaria* (Figs 3b, 4, S1). By expanding taxon sampling of the New World *Eutelia* (from Costa Rica, Bolivia, USA, and French Guiana) in our COI + multi-locus phylogeny (Figs 4, S1), we found that they were all nested within the Nearctic *Marathyssa*, corroborating the results of the multi-locus phylogeny. As a result, the New World species of *Eutelia* (e.g., *E. pyrastis* and *E. pulcherrimus*) should be transferred to *Marathyssa* (Table 5). *Eutelia cuneades + Marathyssa cuneata* (Fig. 4, S1) together form the sister-group of *Penicillaria* and require further morphological study (Table 5). *Phlegetonia violescens* Hampson appears to be closely related to the *Targallodes* species group (Figs 1, S1), and the new combination *Targallodes violescens* **comb nov**. is implemented here (Table 5). Furthermore, a detailed investigation of the original description and barcode analysis of the African species, *Eutelia megacycla* Berio, revealed that this taxon should be synonymized with *Targallodes violescens*. The TS of the African genus *Phlegetonia* (*catephioides* Guenée) is grouped within the true *Eutelia* clade (Fig. S1). Thus *Phlegetonia* should be synonymized with *Eutelia* as suggested by Holloway (1985), although the name has been used in many publications and online databases, such as Afromoths (De Prins and De Prins, 2011). Holloway (1985) concluded that the male and female genitalia of *catephioides* are of the same general form as those of the type species of *Eutelia*, the Mediterranean *E. adulatrix*. Therefore, *Phlegetonia* is once again here placed as a junior subjective synonym of *Eutelia* (Table 5). Our time of divergence analysis reveals that the initial split between the Old World *Eutelia* + the Afro-Oriental *Targallodes* and the Nearctic *Marathyssa* began in the late Oligocene (∼ 25.47 my), resulting in the trans-Atlantic split into two distinct lineages (*Eutelia + Targallodes* and *Marathyssa*) (Fig. 5). Following this, the *Eutelia + Targallodes* lineage diverged between the Old World lineage (*Eutelia*) and the Afro-Oriental lineage (*Targallodes*) (∼ 22.49 my).

*Mimanuga* + *Anuga canescens* (Walker) are together recovered as the sister to the *Anuga* clade + *Phalga*. Poole (1989) placed *Mimanuga* as a synonym of *Anuga* but our results support the monophyly of *Anuga* (excluding *Anuga canescens*) and a close relationship between *Mimanuga* and *Anuga canescens* (Fig. S1). As a result, we reinstate *Mimanuga* from synonymy and transfer *Anuga canescens* into *Mimanuga* as *Mimanuga canescens* **comb. nov**.

Our coverage of New World taxa included all five genera previously assigned to the family in this geographic realm (Poole 1989). Neotropical Euteliidae are comparatively few, the majority comprising a moderately large diversification of *Paectes* with about 30 described and at least as many undescribed species. The Neotropical genus *Thyriodes* has been included in Euteliidae (Poole 1989), but examination of the larval morphology and hostplant (Janzen website), in addition to adult morphology, suggested that this genus belongs to the Erebinae. Our results corroborate a close relationship of *Thyriodes* with members of the subfamily Erebinae within Erebidae (Fig. 4), and we accordingly transfer *Thyriodes* to Erebidae: Erebinae. Tribal placement within Erebinae will require greater coverage of taxa and gene sampling. Stictoperinae are very poorly represented in the Neotropics, *Nagara* being the only genus confirmed there. Thus, only three unrelated and relatively small lineages of Euteliidae occur in the New World: *Nagara* (Stictopterinae: Stictopterini), *Marathyssa* (= *<u>Eutelia</u>*) (Euteliinae), and *Paectes* (*Paectes*) (Euteliinae). The biogeography and ancestral occurrence of these lineages remain untested, but Neotropical colonisations from Old World ancestors certainly warrant consideration.

*Aplotelia diapera* is placed between the remainder of *Aplotelia* and *Chlumetia* (Fig. 4). Although we would note that it is also treated as *Eutelia diapera* in numerous resources (Poole 1989; Markku Savela 2022 https://ftp.funet.fi/pub/sci/bio/life/insecta/lepidoptera/ditrysia/noctuoidea/euteliidae/), our results suggest the species requires morphological study and potentially a new genus.

The two tribes of Stictopterinae identified in the previous studies (Holloway, 2011) as Stictopterini and Odontini were recovered here as monophyletic groups with good support (Fig. 3). *Odontodes, Lophoptera* (both represented by TS), *Sigmuncus, Savoca* and a singleton *Diascoides* with only COI data are the main genera of the tribe Odontodini (Fig. 4). *Lophoptera* is found to be polyphyletic since the Southeast Asian species *Lophoptera huma* is not associated with other *Lophoptera* (including the TS) and needs to be included in a new genus that is more likely associated with the *Gyrtona* clade (Figs 3b, 4, S1). Below, we describe a new genus to accommodate it, as it is evidently distinct from *Lophoptera* species-group.

Holloway (1985) found it difficult to establish clear morphological differences between *Odontodes* and *Lophoptera*, but based on morphological features of male and female genitalia, he established a system that recognized a series of groupings. However, many of these were monobasic or with just two species. Our analysis supports the distinction of the two genera and also supports two of Holloway’s larger groups within *Lophoptera*. His groups B1 and B6 fall into our clade from *L. stipata* (Walker) to *L. nama* (Swinhoe) (Fig. S1) that contains the type-species of the synonym genus *Evia* Walker, *E. ferrinalis* Walker. His group D1 is recovered as our clade from *L. melanesigera* Holloway to *L. squammlinea* Holloway, and includes *L. squammigera* Guenée, the type-species of *Lophoptera* (Fig. S1). In contrast, we found (Fig. 4, S1) that the genus *Sigmuncus* falls into a clade from *L. brunnistis* Holloway to *L. arcuata* (Hampson), the latter being the second species included in *Sigmuncus* by Holloway (1985) besides the here unsampled type species, *S. albigrisea* (Warren). Most of these species have a modification like a twist or scroll to the apex of a slender uncus, which can be treated as a synapomorphy for the whole clade that brings together species from groups A1 and D5, and thus *Sigmuncus* is better treated as a synonym: *Lophoptera* = *Sigmuncus* **syn. nov**.

Like *Sigmuncus*, the sampled members of the genus *Savoca* are nested within *Lophoptera* on both nDNA and barcode data (Figs 4, S1). This is a group of small species that were distributed amongst the genera *Lophoptera* and *Gyrtona* in previous classifications (Holloway, 1985). The species are relatively small, with rectangular forewings patterned usually with a distinctive basal zone and a postmedial that is fine, double, biarcuate, roughly perpendicular to the dorsum. The hindwings often grade paler based but are not hyaline. Hence *Savoca* is here treated as a synonym: *Lophoptera* = *Savoca* **syn. nov**.

The sister tribe, Stictopterini, includes the genera *Stictoptera* (including the TS), *Aegilia* (including the TS), *Gyrtona*, the new genus described below for *Lophoptera huma*, and the Neotropical genus *Nagara* (including the TS).

### *Niklastelia* description

*Niklastelia* Zahiri & Holloway **gen. nov**.

**Type species**: *Eutelia apicifascia* Hampson, 1894(Figs 7a–f)

**Fig. 7.**
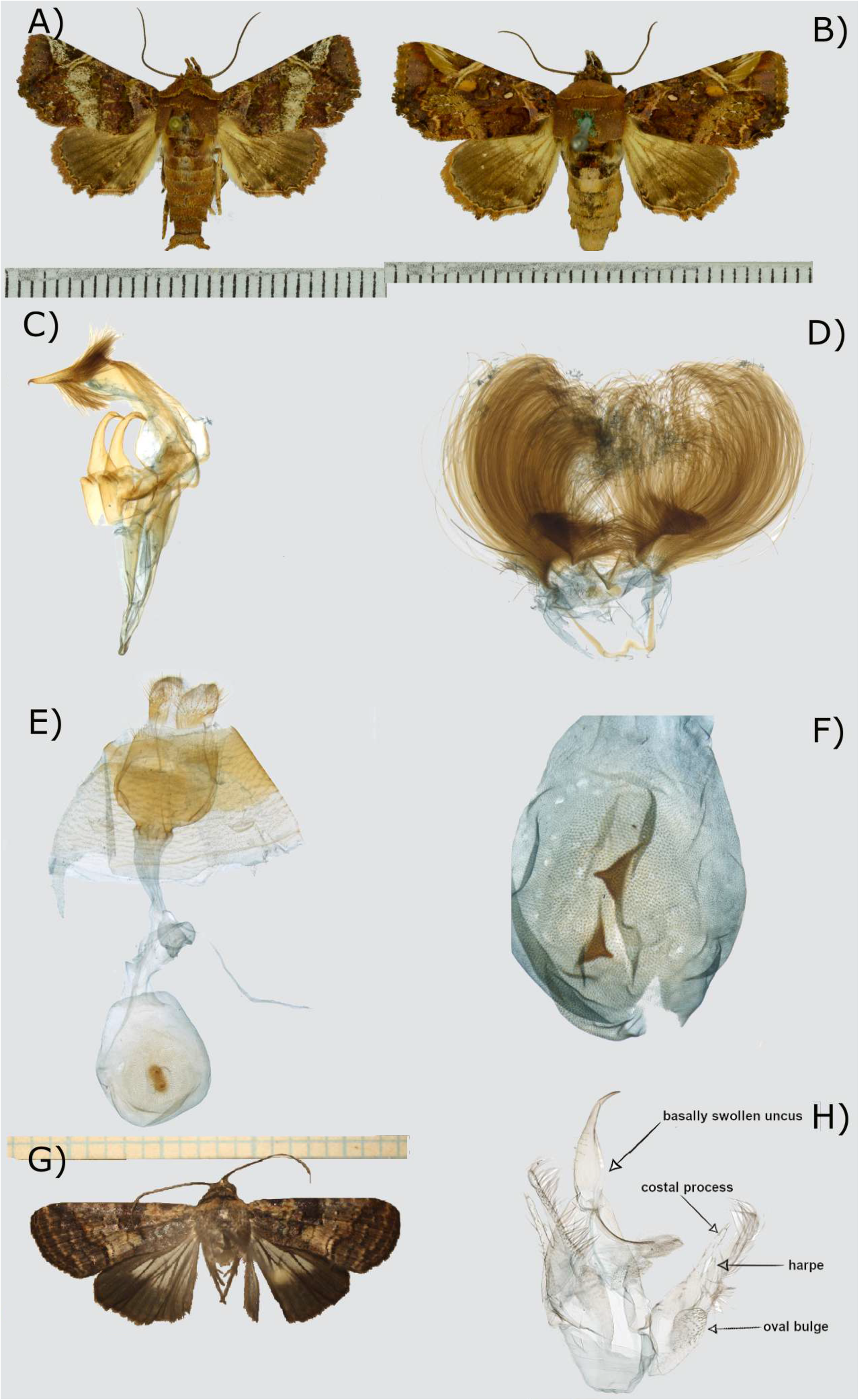
Photos of the new genera *Niklastelia* and *Pellinentelia*: **A)** adult male of *Niklastelia apicifascia* (coll. RMNH) (photo by Rob de Vos), Indonesia, Papua Kab, Yahukimo, WALMAK (distr. Nipsan), 4° 07’ S – 138° 36’ E // 1710 m. at light, 20-25.ix.2011, leg. F. Groenen // RMNH.INS.1557720; **B)** adult female of *Niklastelia apicifascia* (coll. RMNH) (photo by Rob de Vos), Fort de Kock (Sumatra), 920m., November 1921, leg. E. Jacobson // RMNH.INS.1557723; **C)** male genitalia of *Niklastelia apicifascia* (RMNH.INS.1557722) (dissection and photo by Rob de Vos); **D)** eighth abdominal segment of male of *Niklastelia apicifascia* with coremata (RMNH.INS.1557722) (dissection and photo by Rob de Vos); **E)** female genitalia of *Niklastelia apicifascia* (RMNH.INS.1557721) (dissection and photo by Rob de Vos); **F)** bursa copulatrix of female genitalia of *Niklastelia apicifascia* with a pair of scobinate signa (RMNH.INS.1557723) (dissection and photo by Rob de Vos); **G)** *Pellinentelia huma* from Thailand (photo by M. Pellinen); **H)** male genitalia of *Pellinentelia huma* from Thailand (dissection and photo by M. Pellinen).

**LSID**: urn:lsid:zoobank.org:act:22F2BCA5-92F7-439C-83FB-968810657353

The type species has been placed variously in *Eutelia, Phlegetonia* and *Targalla*, but was indicated to be somewhat atypical for all of these by Holloway (1985). The male antennae are weakly bipectinate basally (filiform in typical *Targalla*). The male genitalia are distinctive with valves reduced, slender, strongly upcurved at the apex (Fig. 7c). The genus lacks the marked flexing of the tegumen of the male seen in typical *Targalla*. The eighth abdominal segment bears coremata as in other *Targalla* (Fig. 7d). In the female, the bursa copulatrix has a pair of scobinate signa rather than several patches of some weak scobination basally (Figs 7e, 7f).

Our analysis places the genus in a position well separated from typical *Targalla* or its subgenus *Euteliella* Roepke, but associates it instead with *Paectes* and *Atacira*, further support for its generic status.

The genus consists of the type species, *Niklastelia apicifascia* **comb. nov**., which occurs throughout the Indo-Australian tropics as far east as the Solomon Islands, and *Niklastelia barbara* (Robinson) **comb. nov**. from Fiji and Samoa.

#### Etymology

the genus is named for Niklas Wahlberg in recognition of his work on the phylogeny of the Noctuoidea.

### *Pellinentelia* description

*Pellinentelia* Holloway & Zahiri **gen. nov**.

**Type species**: *Stictoptera huma* Swinhoe, 1903(Figs 7g, 7h)

**LSID:** urn:lsid:zoobank.org:act:A2CC2497-A688-4D7D-88F2-A7E96E1951AA

The type species, *Stictoptera huma*, was originally placed in *Stictoptera* by Swinhoe (1903) and then transferred to *Lophoptera* by Hampson (Hampson, 1912). Holloway (1985) placed it in his “D group” of *Lophoptera*, which was characterized by a synapomorphy of the female genitalia: the presence of both the post-ostial invagination and lateral lobes. It was placed as the sole species in subgroup “D2”. The post-ostial invagination is short, triangular, and the lateral lobes are small semicircular flaps. The male genitalia have a basally swollen uncus, a rod-like costal process almost as long as the valve, and a small, centrally placed harpe (Fig. 7h). These last two features only came into focus when our analysis placed the species well within the Stictopterini and not in the Odontodini, as they represent synapomorphies that are masked by the enlargement of the sacculus.

*Pellinentelia huma* **comb nov**., the only included species, is distinguished from the smaller *Lophoptera* and *Gyrtona* by its robustness, the marked basal hyaline area of the hindwing, and the presence of two black patches on the costal margin of the forewing (Fig. 7g). In the male genitalia, the sacculus bears an oval bulge, with a process on the apex.

As indicated earlier, our analyses placed the genus in a position well separated from typical *Lophoptera*, but in a sister relationship with *Gyrtona*, further support for its generic status.

The genus occurs throughout Thailand, Sumatra, Borneo, Sulawesi, Seram, and New Guinea (Holloway 1985), and China (Hainan, Guangxi) (Qi et al., 2011).

#### Etymology

The genus is named for Markku Pellinen in recognition of his work on the Moths of Thailand.

### Hostplant associations and divergence time of major lineages of Euteliidae

The plant family Anacardiaceae (order Sapindales) provides the majority of the larval hostplants of Euteliinae (Fig. 6). Other major hostplant families are also lactiferous, possibly indicating some degree of preadaptation for host-shifts of Odontodini from Clusiaceae (order Malpighiales) to exploit Dipterocarpaceae (order Malvales). Dipterocarpaceae are a highly diverse family in Southeast Asia that are a significant component of the plant foliage biomass yet have a low load of defoliators (Holloway, 1989). For example, in Bornean forests, dipterocarps are represented in most moist lowland forest types, with ten genera and at least 267 species, and usually about three-quarters of large trees in Bornean lowland forest are dipterocarps (Whitmore, 1984). This dominance is thought to have developed from the Middle Miocene (12–14 my) in response to climatic shifts (Cvetković et al., 2022). Dipterocarps, unlike the majority of flowering plants, are resinous like gymnosperms. In temperate forests where gymnosperms are common, the most significant divergence in feeding preferences in Macroheterocera is between angiosperms and gymnosperms, with much lower diversity of the moths that feed on them (Holloway and Hebert, 1979) in the latter. However, this prominence of gymnosperms gives way to an almost total predominance of angiosperms in subtropical and tropical forests. As a result, it could be speculated that dipterocarps occupy the “conifer niche” in Southeast Asian lowland forests in the absence of gymnosperms. In temperate forests, only highly polyphagous species such as in the erebid subfamily Lymantriinae and the geometrid subfamily Ennominae appear able to span this discontinuity and feed on both angiosperms and gymnosperms (Fig. 8). It is therefore perhaps not surprising that very few Lepidoptera made the switch to dipterocarps and are now specialists or even predominant feeders of the family. The high species richness of Odontodini in these habitats might thus be explained by their exclusive specialism on dipterocarp hostplants.

**Fig. 8.**
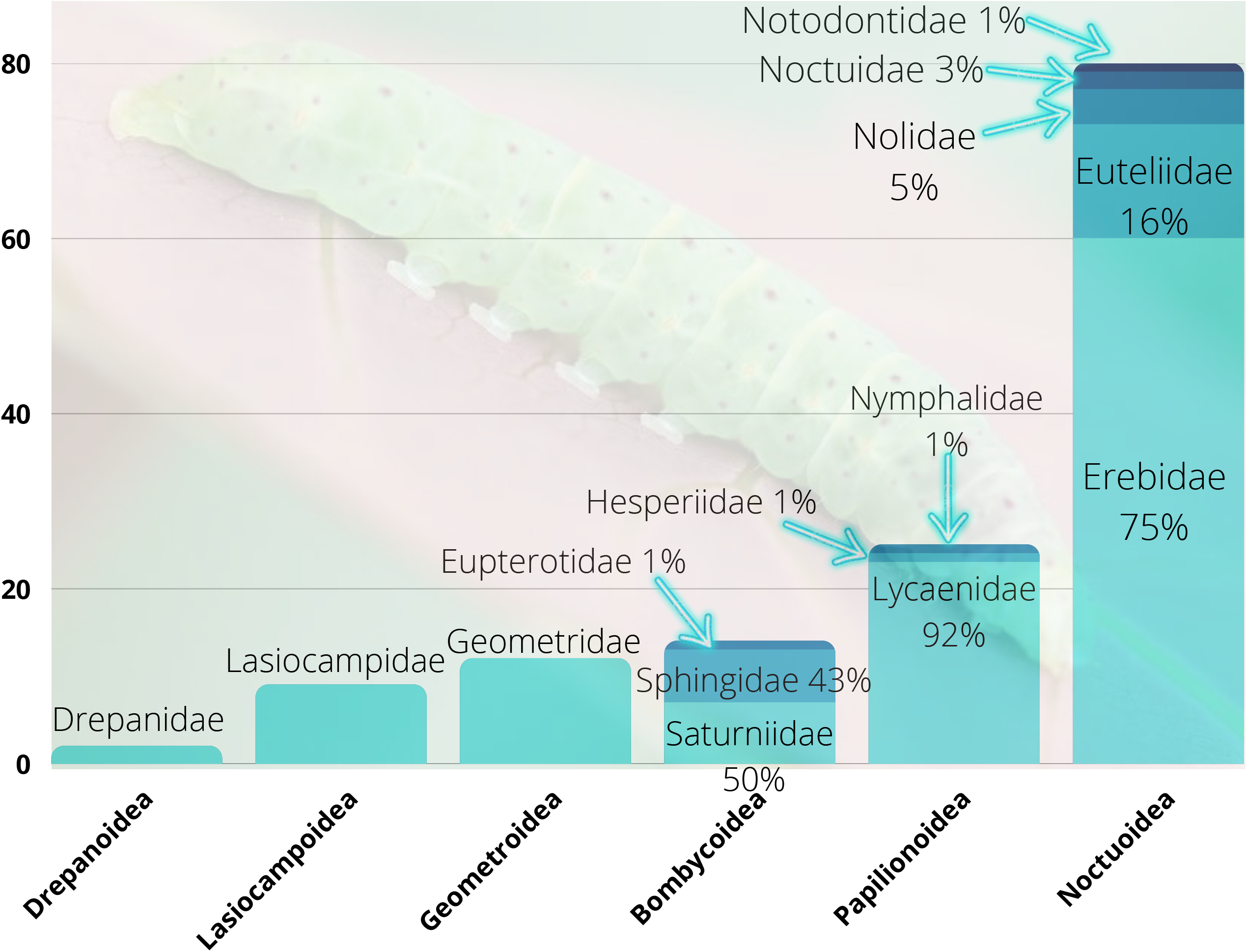
Lepidopteran lineages feeding on the Dipterocarpaceae, the dominant family in most moist lowland forest types of the Oriental tropics.

Our ancestral trait reconstruction identified the plant order Malpighiales (e.g., Clusiaceae and Euphorbiaceae) as the ancestral hostplant for the subfamily Stictopterinae and the tribe Stictopterini. The sister lineage (Odontodini) could have switched from Malpighiales to Malvales (e.g., Dipterocarpaceae) once they became available as a resource. Our time of divergence analysis also revealed that the diversification of major generic lineages of Stictopterinae appears to have begun approximately in Middle Miocene (12–14 my). These estimates are similar to those recovered in phylogenomic divergence time estimations of Dipterocarpaceae (Cvetković et al., 2022), which suggested a species diversification from the Late Oligocene–Early Miocene (21–27 my) in most lineages except for Richetioides (Middle Miocene 14.05 my). The reconstruction of ancestral character states for host-plants pairs several genera with Fabales. However, this maybe not be a positive switch because the Fabales come out at the very top of Holloway’s table for plant orders across the Macroheterocera, and so the switch may just be a reversal to an ancestral state (i.e., plesiomorphy). The divergence of the *Atacira* clade was estimated at the Miocene/Pliocene (∼ 7 my) boundary. Most are lowland species in Borneo, and the host range is diverse within the usual euteliid range (no dipterocarps recorded). The genus certainly occurs in dipterocarp forests but may prefer alluvial and swamp forest systems.

The situation in Lymantriinae (Erebidae) has been studied in depth by Wang et al. 2017), with feeding on Dipterocarpaceae evident in several lineages, both large and small. Using the Oriental tropical data from Robinson et al. (2001) as processed by Holloway (unpublished), but summarized by Holloway (2019), the scores for Lymantriinae overall were 180 for Fabaceae. Then, leaving aside Poaceae with several specialist lineages, the next four plant families all have resin or latex in their sap, totaling together a score of 202: 62 Euphorbiaceae, 51 Dipterocarpaceae, 47 Anacardiaceae and 42 Myrtaceae (Fig. 8). The three most diverse internal lymantriine clades all have high scores for Dipterocarpaceae: 13 in Lymantriini (all in *Lymantria* Hübner), 18 in Orgyiini, and 10 in Nygmiini (Fig. 8). However, feeding on Dipterocarpaceae is predominant at 10 in the small tribe Arctornithini, which is the second most basal branch of Lymantriinae phylogeny as a whole. This tribe consists solely of the genus *Arctornis* Germar, which has numerous species in Southeast Asia of which no fewer than 78 have been recorded so far in Borneo (Holloway, 2011). The most basal branch is Daplasini, a monobasic tribe without host records. The next tribe from the base, the Leucomini, has one genus, *Perina* Walker that is a specialist on *Ficus* L. (Moraceae, also with latex). It may be that polyphagy dwindles with the age of a clade leaving it restricted to the host or hosts to which it became, in some way, uniquely adapted.

In the basal family of Noctuoidea, the mainly arboreal-feeding Notodontidae, there is only one record on Dipterocarpaceae, from *Brykia* Gaede in subfamily Ceirinae. In the four quadrifid families only the Erebidae, with 60 records, and the Euteliidae, with 13, show any significant exploitation of dipterocarp foliage, each showing some degree of specialism (Fig. 8). There are three records in Nolidae, two from the polyphagous and generally arboreal subfamily Nolinae, and one from Eligminae where records from more temperate latitudes and a few postdating Robinson et al. (2001) indicate the possibility of a minor degree of specialism associated with feeding also on Fagaceae (Symondson et al., 2015), as in the lycaenid genus *Arhopala* Boisduval discussed later. The only two records in Noctuidae are for the highly polyphagous genus *Spodoptera* Guenée in Prodeniini, although Holloway (2011) noted a fruit-feeding record for *Elusa* Walker, a genus of uncertain placement.

In Euteliidae, dipterocarp feeding predominates in the stictopterine tribe Odontodini with eight records, followed by four for Euphorbiaceae. The sister-tribe, Stictopterini, has one record, possibly an error through past confusion of generic concepts, as most records (12) are from the Clusiaceae. Euteliinae have four records, though this subfamily feeds predominantly on Anacardiaceae (32 records) with a subgroup of *Targalla* and the genus *Pataeta* on Myrtaceae. Thus, no clear instance in the Macroheterocera of exclusive specialism at a generic level has been discovered, and genera with a high incidence of dipterocarp feeding are few and far between, but three of those genera are among the more species-rich in the region: *Arhopala* in Lycaenidae; the related genera *Lophoptera* and *Odontodes* (tribe Odontodini of the Stictopterinae) in Euteliidae; and *Arctornis* in Lymantriinae. Two further genera in Geometridae, *Ornithospila* Warren in Geometrinae and *Ectropidia* Warren in Ennominae, each have a unique record from Dipterocarpaceae. This is not significant in itself but species of both genera have been found to decline in numbers or be absent in samples from forests where dipterocarps had been logged out compared with undisturbed forests of the same type (Intachat et al., 1999), and five species of *Ornithospila* were almost exclusive to a sample from an indigenous plantation consisting mainly of dipterocarps compared with those from three other areas that had undergone various intensities of logging (Intachat et al., 1997). These observations were made in Peninsular Malaysia. Thus, the three genera most successful at exploiting the dipterocarps seem to have achieved this by slightly different evolutionary routes. Though none of the genera is exclusively specialist on dipterocarps, all three are species-rich in SE Asia, particularly Borneo, where current lists have 48 species for *Lophoptera*, 77 for *Arctornis* and 89 for *Arhopala* (Seki et al., 1991).

## Conclusions

Despite the rather weak support for deep internal nodes and short branches that are not well supported, we consider that our phylogenetic hypothesis is generally firm enough to be informative of a stable classification for Euteliidae. As we found previously (Zahiri et al., 2011), the results of the current study continue strongly to support the monophyly of four quadrifid noctuoid families. For Euteliidae, we have been able to resolve the relationships of major lineages within the family and we have elucidated the phylogenetic positions of many previously unplaced taxa. Furthermore, DNA barcodes combined with multigene backbone DNA data, as previously demonstrated in the subtribe Polyommatina (Lepidoptera: Lycaenidae), a large lineage including numerous species with unresolved taxonomy (Talavera et al., 2022), can be used successfully to generate reliable phylogenetic hypotheses and significantly improve higher-level systematics in large datasets.

The diversification of phytophagous insects is likely associated with their tendency to specialize on specific host plants. This specialization may induce host shifts with numerous expansions and reductions of the host-plant range (Janz et al., 2001) that eventually promote the evolution of new specialist species. This study reveals that conservatism in utilizing lactiferous hostplant families is prevalent in the moth family Euteliidae. *Lophoptera* and *Odontodes* are the main genera of the tribe Odontodini with strong larval feeding preferences for species of Dipterocarpaceae. The sister tribe, Stictopterini, appears to be restricted to Clusiaceae, apart from two records from Dipterocarpaceae. In the other subfamily, Euteliinae, Anacardiaceae are predominant as larval hosts. Thus, all hosts in the family are lactiferous, possibly providing some degree of preadaptation for exploiting Dipterocarpaceae. All these plant families have their highest diversity in the Oriental tropics, hence the occurrence of over 30% of known Euteliidae in Borneo is not surprising. The hypothesis that this richness is correlated with the regional concentration of dipterocarps has yet to be tested for *Lophoptera* and *Odontodes. Arctornis* might provide an ideal testing ground, as the species appear to be geographically more localized than those of *Lophoptera*, with an already greater accumulation of host information. Collection of larvae from tree samplings including dipterocarps is producing many more records of dipterocarp-feeding in the genus, as well as an indication of significant diversity of form in the tufts of secondary setae exhibited by the larvae. Barcoding of the genus is building up both for Borneo and New Guinea, but much more sampling and morphological investigation is needed through the range.

## Declarations

### Consent for publication

The manuscript has been approved for submission by all co-authors and is not currently under consideration in any other journals.

All authors read and approved the final manuscript.

### Availability of data and materials

All data generated or analyzed during this study are accessible in GenBank through accession numbers provided in Tables 1 & 3.

### Ethics approval statement

NA

### Zoobank registration

1. ZooBank publication URL: http://zoobank.org/urn:lsid:zoobank.org.pub: C0ABC829-938D-4409-B544-BB90F69D55EA
2. ZooBank nomenclatural act registration code: ***Niklastelia*:** http://zoobank.org/urn:lsid:zoobank.org:act: 22F2BCA5-92F7-439C-83FB-968810657353 ***Pellinentelia*:** http://zoobank.org/urn:lsid:zoobank.org:act: A2CC2497-A688-4D7D-88F2-A7E96E1951AA

### Authors’ contributions

The following table shows the major contributions of authors to the original article.

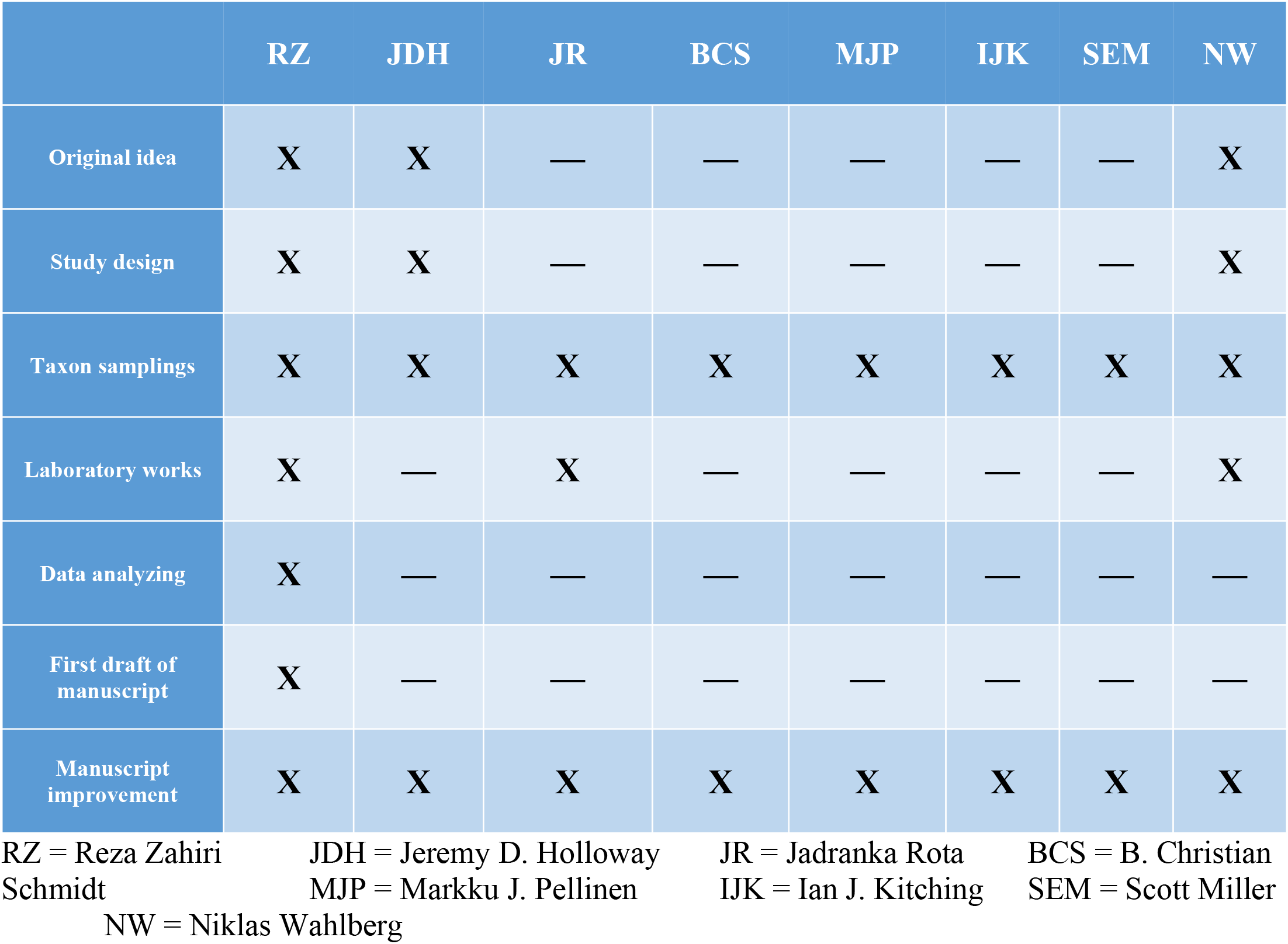

## Acknowledgements

This work was supported financially by the Academy of Finland and the Kone Foundation awarded to N. Wahlberg; CIMO + Finnish Cultural Foundation + Alfred Kordelin Foundation awarded to R. Zahiri; CeNak, University of Hamburg, Germany (supported by Dr. Martin Husemann) to R. Zahiri; Canadian National Collection of Insects, CNC (provided by Chris Schmidt) to R. Zahiri; Canadian Food Inspection Agency, CFIA to R. Zahiri. The main sources of samples, other than those collected by the authors (MJP, BCS, IJK), are the LepTree project, headed by Charles Mitter *et al*. (US NSF award #0531769); the Natural History Museum, London, UK; Daniel H. Janzen (US NSF #DEB0072730 and DEB0515699); Roger C. Kendrick (Kadoorie Farm and Botanic Garden, Hong Kong); Pasi Sihvonen (Helsinki, Finland); Henry Barlow (International Trust for Zoological Nomenclature, UK & Malaysia). The authors also acknowledge Rob de Vos (Naturalis Biodiversity Center, The Netherlands) for the specimen dissection and photography of *Niklastelia apicifascia*. The Papua New Guinea specimens come from a rearing campaign by the Binatang Research Centre led by Vojtech Novotny, George Weiblen, Yves Basset and Scott Miller, and supported by the US National Science Foundation (grant DEB-0211591 and others), Czech Science Foundation grant 206/09/0115 and others, and Czech Ministry of Education & European Union grant CZ.1.-07/2.3.00/20.0064. DNA barcodes were largely provided by Paul Hebert through a grant from Genome Canada to the iBOL project.

## Authors’ information

**Reza Zahiri**:

Canadian Food Inspection Agency, Ottawa Plant Laboratory, Entomology Laboratory, 960 Carling Ave., Ottawa K1A 0C6, Ontario, Canada

**Jeremy D. Holloway**

Science Group, Natural History Museum, Cromwell Road, London SW7 5BD, UK

**Jadranka Rota**:

Biological Museum, Department of Biology, Lund University, Lund, Sweden

**B. Christian Schmidt**:

Agriculture and Agri-Food Canada, Biodiversity Program, Canadian National Collection of Insects, Arachnids, and Nematodes, K.W. Neatby Bldg., 960 Carling Ave., Ottawa K1A 0C6, Ontario, Canada

**Markku J. Pellinen**:

175, M1 Muban Phichai, Mueang Lampang, Lampang, 52000 THAILAND

**Ian J. Kitching**:

Science Group, Natural History Museum, Cromwell Road, London SW7 5BD, UK

**Scott E. Miller**:

National Museum of Natural History, Smithsonian Institution, Washington, DC 20008, USA

**Niklas Wahlberg**:

Department of Biology, Lund University, Lund, Sweden

## Supplementary material captions

**Table S1**. Best models of DNA sequence evolution for the multigene dataset.

**Fig. S1**. Phylogenetic hypothesis of Euteliidae and outgroups inferred by Maximum Likelihood analysis. The MLtree contains 425 terminal taxa in total: 183 with multi-locus sequence data and 242 terminals represented only by mitochondrial COI barcode data. Bootstrap values (please see legend) are displayed as grey circles (five various sizes) on the tree branches. The scale bar shows the estimated number of substitutions per site. The colour bars represent gene regions used in the ML analysis for each leaf: red represents COI-barcode and the rest represent COI3P (mtDNA), EF1a, GAPDH, IDH, MDH, RpS5, Wgl400, CAD (nDNA), respectfully. Type species are marked with red stars.

---

**Table 1**. List of taxa with voucher codes (specimen ID = specimen identity) and GenBank accession numbers used in the multi-locus analysis. The families and subfamilies are classified as indicated in Figs 1 to 6. – = Gene was not amplified for specimens.

**Table 2**. List of generic names with their type species (TS) of Euteliidae.

**Table 3**. List of taxa with voucher codes (specimen ID = specimen identity) and GenBank accession numbers used in multi-locus + COI5P analysis.

**Table 4**. Hostplant data in the family Euteliidae was used in ancestral trait reconstruction using RASP 4.2.

**Table 5**. Taxonomic changes including new combinations, new synonymies, and new taxa.

