## Supplementary figures and images for "Evolutionary history of Euteliidae (Lepidoptera, Noctuoidea)"

### Fig. S1

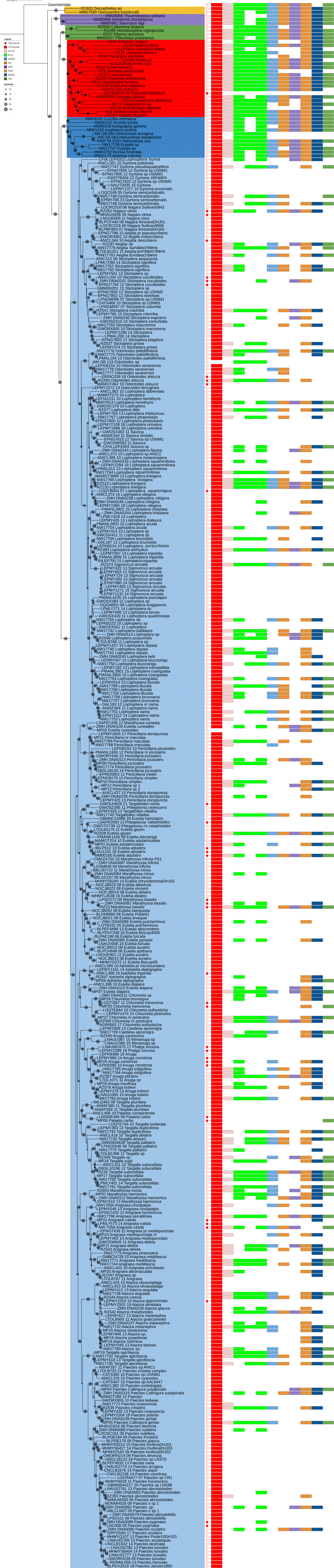
